# mTOR S-nitrosylation inhibits autophagy and lysosomal proteolysis

**DOI:** 10.1101/2020.09.11.292607

**Authors:** Bryce W.Q. Tan, Sijie Tan, Byorn W.L. Tan, Sheeja Navakkode, Cheng Yang Ng, Steven Yuan, Mui Cheng Liang, Chao Liu, Shi Yin, Chou Chai, Katherine C.M. Chew, Yee Kit Tai, Sreedharan Sajikumar, Yulin Lam, Ping Liao, Han-Ming Shen, Kah-Leong Lim, Esther Wong, Tuck Wah Soong

**Affiliations:** Department of Physiology, National University of Singapore, Singapore 117593; Department of Medicine, National University Health System, Singapore; School of Biological Sciences, Nanyang Technological University, Singapore 637551; National Neuroscience Institute, Singapore 308433; Department of Chemistry, National University of Singapore, 3 Science Drive 3, Singapore 117543; Jiangsu Province Key Laboratory of Anesthesiology; Jiangsu Province Key Laboratory of Anesthesia and Analgesia Application Technology, Xuzhou Medical University, Xuzhou, China; Centre for Healthy Ageing, National University Health System, Singapore 117456; LSI Neurobiology Programme & MSC Neuroscience Programme, National University of Singapore 117456; NUS Graduate School for Integrative Sciences and Engineering, National University of Singapore 117456

**Keywords:** Nitric oxide synthase, nitrosylation, mTOR, autophagy, lysosomes, phosphoinositide, Alzheimer’s Disease

## Abstract

Mammalian Target of Rapamycin (mTOR) is a master regulator of autophagy and lysosomes, and its downstream kinase-dependent pathways have been extensively characterized. Here, we report an unexpected kinase-independent regulation of autophagy and lysosomes by S-nitrosylation at Cys423 position of mTOR that resulted in suppression of VPS34 and PIKfyve-dependent phosphoinositide synthesis. Physiologically, S-nitrosylation of mTOR reduced basal lysosomal proteolysis via nitric oxide synthase (NOS)-mediated synthesis of NO from lysosomal arginine precursor, a marker of cellular nutrition status. Significantly, we found increased lysosomal NOS-mTOR complexes in APP-PS1 Alzheimer’s disease (AD) murine model, and increased mTOR S-nitrosylation in AD patient-derived fibroblasts. Lastly, we demonstrated that pharmacological inhibition of NOS or overexpression of mTOR^Cys423Ala^ mutant reversed lysosomal and autophagic dysfunction in AD patient-derived fibroblasts, suggesting novel therapeutic strategies for autophagosome-lysosomal activation.

## Introduction

S-nitrosylation of proteins, including XIAP(*1*) and PDI(*2*), plays a critical role in neurodegeneration(*3, 4*). While recent evidences suggest that arginine is a critical amino acid in mTOR regulation of lysosomal activity(*5, 6*), the pathophysiological relevance of lysosomal arginine is not well understood. Emerging studies have implicated aberrant S-nitrosylation(*7, 8*) and dysfunctional arginine metabolism(*9, 10*) in Alzheimer’s Disease(AD) where autophagosome-lysosomal failure contribute to tau and amyloid burden(*11-13*). Taken together, these studies suggest a possible pathophysiological link between arginine, nitrosative stress and autophagosome-lysosomal failure. As arginine is important for nitric oxide synthesis, and the physiological role of lysosomal nitric oxide synthase (NOS) is poorly understood(*14*), we hypothesize that lysosomal arginine, under nutrient-replete conditions, serves as a critical substrate of lysosomal NOS that mediate mTOR S-nitrosylation. This physiological cellular mechanism to sense lysosomal arginine stores will lead to appropriate downstream inhibition of autophagy, while excessive mTOR S-nitrosylation under pathophysiological conditions such as AD contribute towards autophagosome-lysosomal dysfunction.

## Results

### NO mediates autolysosome dysfunction

To determine the effect of NO on autophagy and lysosome function, HEK 293 cells treated with low-dose (4 µM) long-lived NO donor NOC18 have reduced number of autolysosomes **(Figure 1A-B)**, and accumulation of perinuclear lysosomes as detected by lysomotropic dye Lysotracker Red and GFP-LAMP1 puncta **(Figure 1C-D)**, with decreased lysosomal Cathepsin B proteolysis activity**(Figure 1E-F)**. We next investigated effects of NOC18 on other endosomal compartments and did not observe any significant difference in numbers of early endosomes (EEA1-positive; **Figure 1G**), pre-autophagosomes(ATG16L1-positive; **Figure 1H**), or late endosomes (Rab7A-positive; **Figure 1I-J**). Following, we examined if the observed autophagic and lysosomal deficits could be explained by transcriptional dysregulation, and found no alterations in nuclear translocation or phosphorylation of transcriptional factor TFEB (**Figure 1-figure supplement 1 A-D**). By contrast, we found marked TFEB nuclear translocation with high-dose NOC18(500 µM), as was reported by others(*15*). In view of concomitant lysosomal and autophagic deficits with NOC18, we wondered if S-nitrosylation of mTOR, a major signalling regulator of the autophagosome-lysosomal pathway, could be a possible underlying mechanism. Significantly, mTOR was S-nitrosylated, but none of its phosphorylated forms (S2448 or S2481) could be detected, suggesting that only non-phosphorylated forms can be S-nitrosylated (**Figure 1K**). Intriguingly, we observed that mTOR and its substrate p70S6K, ULK1, and AKT phosphorylation levels were not altered with NOC18 (**Figure 1L-O; Figure 1-figure supplement 1 E-G**). In contrast, treatment of Torin-1 reversed NOC18-mediated lysosomal accumulation (**Figure 1-figure supplement 1 H**), impairment of lysosomal proteolysis and autophagy (**Figure 1-figure supplement 1 I-J)**, suggesting kinase-independent effects of S-nitrosylated mTOR. This corroborates with initial observations that only non-phosphorylated forms of mTOR could be S-nitrosylated. As VPS34 and PIKfyve are two major kinases regulating autophagy and lysosomal activity, we next hypothesized if mTOR functionally regulate these kinases in NOC18-treated cells. Of note, there were increased mTOR-VPS34 and mTOR-PIKfyve interactions in NOC18-treated cells (**Figure 1P**) that were rapamycin-insensitive(**Figure 1Q**), while no significant increase in Rictor or Raptor binding with mTOR was observed with NOC18 treatment(**Figure 1P**). These evidences are in keeping with the kinase-independent effects of S-nitrosylated mTOR. Finally, we found that phosphoinositide species PI(3)P and PI(3,5)P_2_, products of VPS34 and PIKfyve respectively, were both downregulated in NOC18-treated cells(**Figure 1R**), which is in keeping with previous studies on the regulatory role of mTOR on these kinases(*16, 17*).

**Figure 1:**
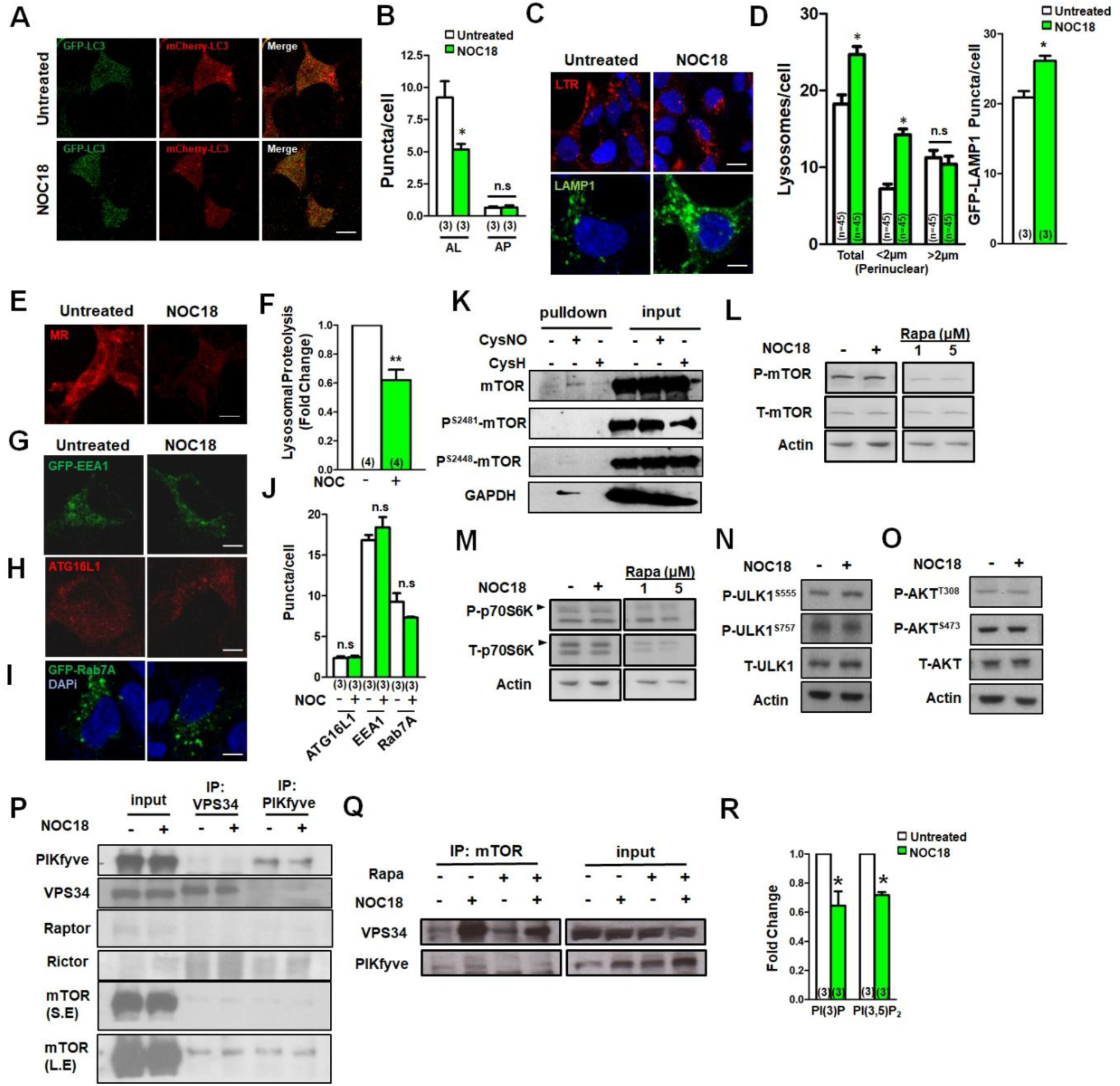
S-nitrosylation of mTOR is kinase-independent. **(A)**mCherry-GFP-LC3-transfected HEK293 cells treated with NOC18 and **(B)**quantification of autolysosomes (AL) and autophagosomes (AP). **(C)**Lysotracker staining and GFP-LAMP1 overexpression of NOC18-treated HEK293 cells and **(D)**quantification of lysosomal distribution and GFP-LAMP1 puncta. **(E)**Magic Red Cathepsin-B substrate staining of NOC18-treated cells and **(F)**quantification graph. **(G)**GFP-EEA1-, **(H)**mCherry-ATG16L1-, and **(I)**GFP-Rab7A transfected HEK293 cells and **(J)**puncta quantification. **(K)**Resin-assisted capture of mTOR and GAPDH. Immunoblot of **(L)**phosphorylated mTOR, **(M)**p70S6K, **(N)**ULK1 and **(O)**AKT. **(P)** VPS34 and PIKfyve immunoprecipitation, and detection of mTOR, Rictor and Raptor(S.E. – short exposure for input; L.E.–long exposure for immunoprecipitated VPS34 and PIKfyve) **(Q)** mTOR immunoprecipitation and detection of VPS34 and PIKfyve. **(R)**HPLC detection of phosphoinositides in NOC18-treated HEK293 cells. Sample size numbers are expressed in parenthesis, and pooled data is expressed as mean (SEM). *p<0.05;**p<0.01;Scale bar:10µm. **Supplementary data: Figure 1-figure supplement 1**

### NO downregulates phosphoinositides

We proceeded to supplement NOC18-treated cells with PI(3)P, and its isoforms PI(4)P or PI(5)P, to ascertain specificity of PI(3)P effects. We observed that PI(3)P and PI(4)P resulted in increased lysosomes with no further increase in lysosomes with NOC18(**Figure 2A**,**D**), whereas PI(3)P treatment abolished NOC18-mediated inhibition of lysosomal proteolysis(**Figure 2B**,**E**) and decrease in autolysosomes(**Figure 2C**,**F**). Our observations on the attenuation of NOC18 effects on autolysosomes with PI(5)P supplementation(**Figure 2C**,**F**), is also consistent with earlier observations that PI(5)P can activate autophagy in PI(3)P-depleted cells(*18*). Part of this phenomenon could be attributed to an increase in lysosomes with PI(5)P supplementation(**Figure 2A**,**D**). On the other hand, PI(4)P did not activate autophagy, contrary to previous observations(*18*), but supplementation of PI(4)P suggested possible lysosomal proteolytic inhibition. Conversely, no further lysosomal accumulation with NOC18 treatment in HEK 293 cells with siRNA-knockdown of VPS34(**Figure 2-figure supplement 1 A)**, nor was there any further decrease in lysosomal proteolysis or autophagic impairment(**Figure 2-figure supplement 1 B-C**). Overall, our observations suggest that NOC18-mediated reduction of PI(3)P levels via VPS34 inhibition could account for NOC18-mediated lysosomal proteolytic and autophagic inhibition. As PI(3)P is essential for PI(3,5)P_2_ synthesis, we next tested whether VPS34 effects are mainly via PI(3)P or its downstream metabolite PI(3,5)P_2_. We proceeded to knock down ATG14, another component of the VPS34-kinase complex, and supplemented the cells with either PI(3)P or PI(3,5)P_2_. First, we confirmed that ATG14 siRNA knockdown reduced intracellular PI(3)P compartments, and supplementation of PI(3)P in ATG14 siRNA-knockdown cells increased PI(3)P compartments (**Figure 2-figure supplement 1D**). Next, we found that autophagic impairment by ATG14 was reversed with PI(3)P but not PI(3,5)P_2_(**Figure 2-figure supplement 1D**), whereas lysosomal accumulation by ATG14 was attenuated by PI(3,5)P_2_ but not PI(3)P (**Figure 2-figure supplement 1 E**), suggesting that the effects of VPS34 on PI(3)P and PI(3,5)P_2_ synergistically modulate lysosomal accumulation and proteolysis, and autophagy. Finally we validated the knockdown efficacies of both siRNAs (**Figure 2-figure supplement 1 F**,**G**): despite significant levels of hVPS34 persisted after siRNA transfection, we still observed significant autophagic and lysosomal deficits in hVPS34-siRNA transfected cells. Finally, no alterations in the Beclin1-VPS34-ATG14 kinase complex was observed, whereas mTOR interaction with VPS34 is enhanced with NOC18(**Figure 2-figure supplement 1 H**), suggesting that the mTOR-VPS34 complex is unique and does not interfere with Beclin1-VPS34-ATG14 complex interaction.

**Figure 2:**
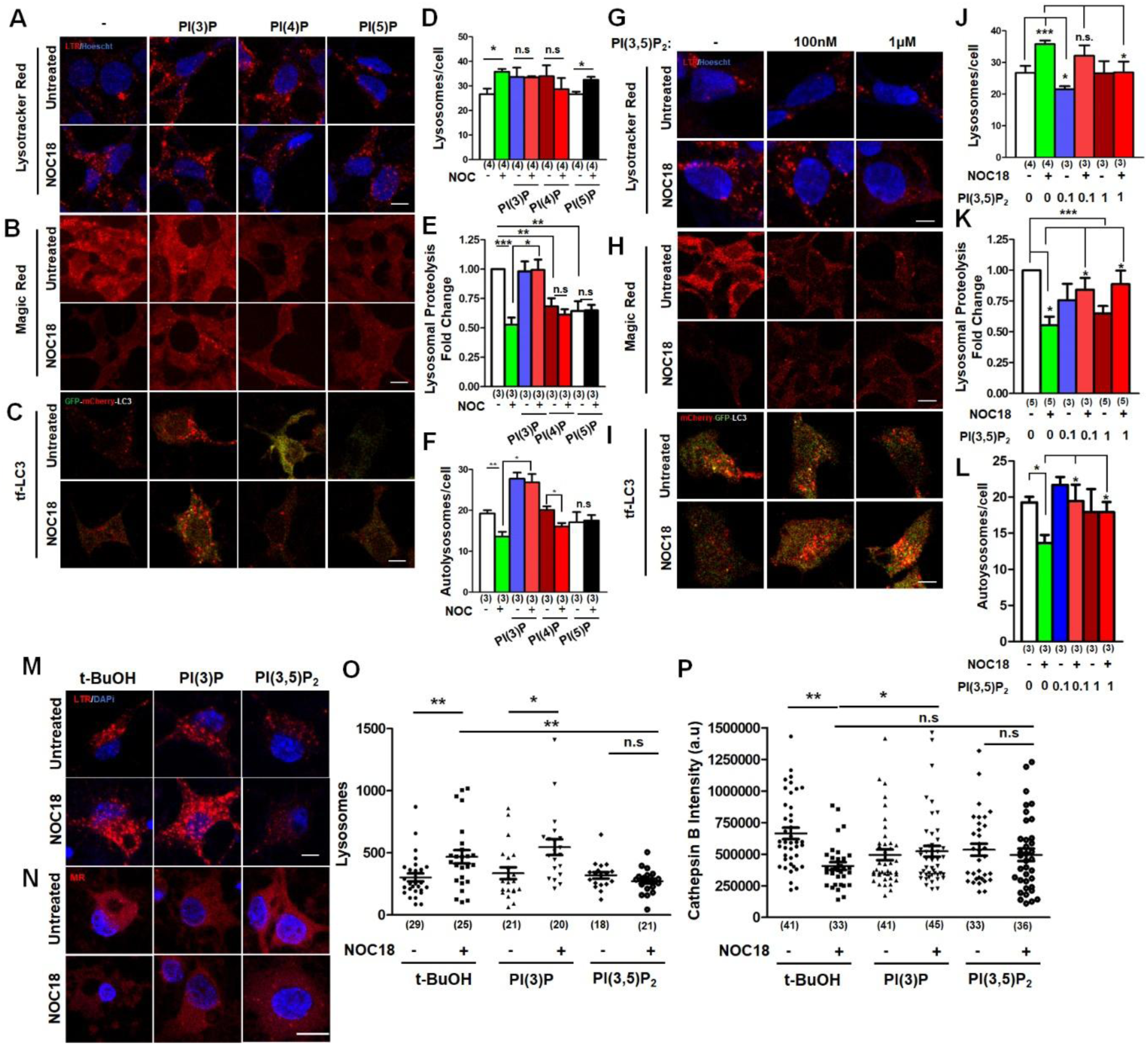
PI(3)P and PI(3,5)P_2_ supplementation reversed NO-mediated inhibition of lysosomal proteolysis. **(A)**Lysotracker Red and **(B)**Magic Red staining, and **(C)**overexpression of mCherry-GFP-LC3 (tf-LC3) in HEK293 cells supplemented with PI(3)P, PI(4)P or PI(5)P, and treated with NOC18, and their respective quantifications**(D-F). (G)**Lysotracker Red staining, **(H)**overexpression of tf-LC3 and **(I)**Magic Red staining in HEK293 cells treated with PIKfyve inhibitor YM201636. **(J)**Lysotracker Red and **(K)**Magic Red staining, and **(L)**overexpression of tf-LC3 in HEK293 cells treated with 100nM or 1µM PI(3,5)P_2_ and NOC18. **(M)** Lysotracker Red and **(N)** Magic Red staining of rat cortical neurons treated with NOC18 in the presence of PI(3)P or PI(3,5)P_2_, and **(O, P)** their respective quantification. Sample size numbers are expressed in parenthesis, and pooled data is expressed as mean (SEM). *p<0.05;**p<0.01;***p<0.001;Scale bar:10µm. **Supplementary data: Figure 2-figure supplement 1**

Next we supplemented NOC18-treated cells with PI(3,5)P_2_ and observed that NOC18-mediated lysosomal accumulation was markedly attenuated by PI(3,5)P_2_ (**Figure 2G,J**), while modest increases in lysosomal proteolysis (**Figure 2H,K**) and autophagy (**Figure 2I,L**) were observed with PI(3,5)P_2_ supplementation. Similarly, supplementation of PI(3,5)P_2_ and PI(3)P in rat cortical neurons reversed NO-mediated lysosomal accumulation (**Figure 2M,O**), and proteolysis (**Figure 2N,P**) respectively. We then hypothesized that disruption of PIKfyve-Fig4-Vac14 complex, an essential complex required for PI(3,5)P_2_ synthesis, by NO could mediate lysosomal impairment. We observed that Fig4-, PIKfyve-, and Vac14-siRNA knockdown alone led to lysosomal accumulation, and no further lysosomal accumulation was observed with NOC18 treatment(**Figure 2-figure supplement 1 I**). Paradoxically, we observed attenuation of NOC18-mediated lysosomal proteolysis(**Figure 2-figure supplement 1 J**) and autophagic impairment(**Figure 2-figure supplement 1 K**) in Fig4-, PIKfyve- and Vac14-siRNA transfected cells. **Figure 2-figure supplement 1L** shows knockdown efficacy of these siRNAs. Similarly, we found that YM201636, a PIKfyve inhibitor, attenuated NOC18-mediated inhibition of autophagy (**Figure 2-figure supplement 1 M**) and lysosomal proteolysis(**Figure 2-figure supplement 1 N**), and lysosomal accumulation (**Figure 2-figure supplement 1 O)**. Taken together, our findings suggest that PI(3)P deficiency from VPS34 inhibition mediates autophagosomal and lysosomal proteolytic impairment, whereas PI(3,5)P_2_ deficiency, contributed by PIKfyve and, to a smaller extent, VPS34 inhibition, mediates lysosomal accumulation by NOC18.

### mTOR S-nitrosylation is implicated in AD

We next asked how mTOR S-nitrosylation occurs? By overexpressing eNOS, iNOS and nNOS, we observed that eNOS, but not iNOS or nNOS, recapitulated fully NOC18 effects on lysosomal accumulation(**Figure 3A,D**), lysosomal proteolysis**(Figure 3B,E)**, and autophagic impairment(**Figure 3C,F**), although decreased lysosomal proteolytic activity was observed in iNOS- and nNOS-transfected cells(**Figure 3B,E**). Correspondingly, these alterations in lysosomal and autophagic activity were reversed by L-arginine analogue L-NAME(**Figure 3A-F**), suggesting that NO mediates these observed effects. Interestingly, we observed strong and weak interaction of eNOS and iNOS with mTOR respectively, while only eNOS overexpression recapitulated our earlier observations that mTOR has increased binding with PIKfyve and VPS34(**Figure 3G-I**). Concomitantly, we observed increased S-nitrosylated mTOR levels in eNOS- and iNOS-overexpressing cells(**Figure 3J,K**). To investigate the physiological relevance of mTOR S-nitrosylation, we examined the APP/PS1 Alzheimer’s disease murine model and observed enhanced mTOR S-nitrosylation in APP/PS1 mouse brains(**Figure 3L**). We proceeded to fractionate wild-type and age-matched APP/PS1 brains and immunoprecipitated mTOR from cytosolic and lysosomal fractions. While no nNOS-mTOR complexes in APP/PS1 lysosomal fractions were observed, there was an overall decrease in lysosomal nNOS. Surprisingly, we observed nNOS-mTOR complexes in the cytosolic fraction in the APP/PS1 brains(**Figure 3M**), which could suggest that other cofactors specific in neuronal cell types could be involved in the complex formation. Next, we found increased eNOS-mTOR complexes in the lysosomal fraction and decreased eNOS-mTOR complexes in the cytosolic fraction of the APP/PS1 brains(**Figure 3M**). However, we did not observe significant increase in iNOS-mTOR complexes in both cytosolic and lysosomal fractions(**Figure 3M**). Lastly, we observed that mTOR-PIKfyve interaction occurs in the lysosomal compartment, whereas mTOR-VPS34 interaction occurs in the cytosolic fraction, suggesting that these two complexes are unique and are spatially separated, and are both increased in APP/PS1 brains(**Figure 3M**). To investigate the physiological significance of increased lysosomal mTOR-NOS in the APP/PS1 brains, we isolated lysosomes from these mouse brains and observed increased NOS activity in APP/PS1 lysosomes, suggesting that these complexes are functional(**Figure 3N**). Correspondingly, we observed lower lysosomal β-glucosidase activity in APP/PS1 lysosomes(**Figure 3O**). As lysosomal arginine is a potential substrate for lysosomal NOS, we hypothesize that S-nitrosylation of mTOR could be an indicator for physiological sensing of lysosomal arginine. First, we observed that lysosomal and autophagic activation by EBSS was attenuated with L-arginine supplementation, but not L-NAME(**Figure 3-figure supplement 1 A-B**). Next, we observed EBSS starvation decreased mTOR S-nitrosylation levels, and was rescued by L-arginine supplementation but not L-NAME(**Figure 3P-Q**). In contrast, in rat brain-derived lysosomes, pre-loading lysosomes with L-arginine, but not L-NAME, increased lysosomal NOS independent of mTOR inhibition by Torin-1(**Figure 3R**). Lastly, lysosomes isolated from ATG5-null mouse embryonic fibroblasts have increased NO activity, which suggest that lysosomal NOS is dependent on cellular autophagic flux(**Figure 3S**). Taken together, our data suggests that firstly, lysosomal arginine, a marker of amino acid nutrient status, mediates mTOR S-nitrosylation under nutrient-replete conditions and acts as a physiological regulator of lysosomal proteolysis and autophagy independent of mTORC1. Secondly, mTOR-eNOS lysosomal localization could play a role in mediating mTOR S-nitrosylation and lysosomal NO production from lysosomal L-arginine in AD.

**Figure 3:**
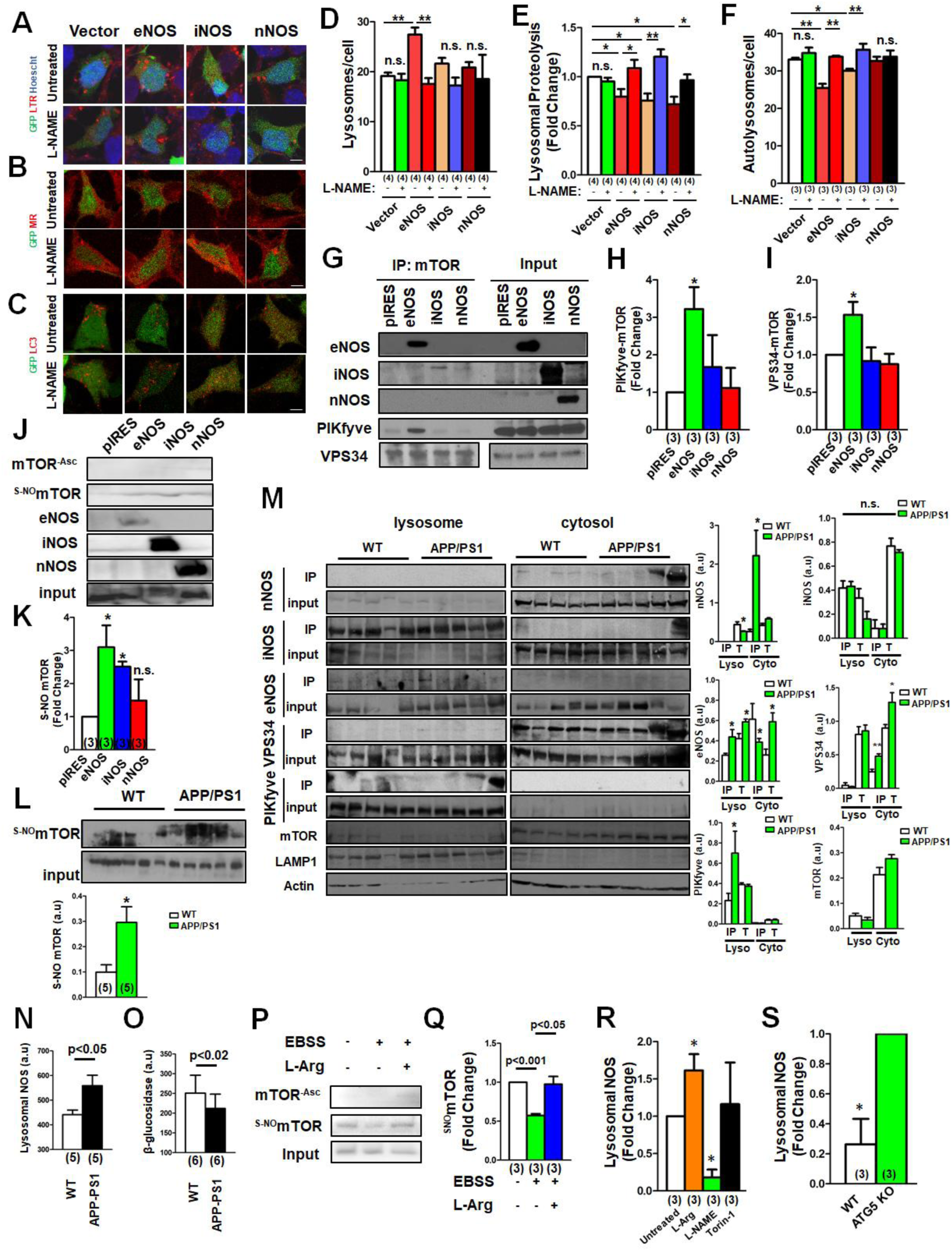
Lysosomal NOS mediates mTOR S-nitrosylation in AD. **(A)**Lysotracker Red and **(B)**Magic Red staining of HEK293 transfected with GFP-pIRES (Vector), GFP-eNOS, GFP-iNOS and GFP-nNOS, and treated with L-NAME. **(C)**Co-transfection of RFP-LC3 with GFP-pIRES, GFP-eNOS, GFP-iNOS and GFP-nNOS, and **(D-F)**their respective puncta quantification. **(G)**Immunoprecipitation of mTOR with overexpression of GFP-pIRES, eNOS, iNOS and nNOS, and immunoblotting for eNOS, iNOS, nNOS, hVPS34 and PIKfyve with **(H-I)** its respective densitometry analysis. **(J)**Resin-assisted capture (RAC) of nitrosylated endogenous mTOR with GFP-pIRES, eNOS, iNOS and nNOS overexpression, with **(K)**its respective densitometry analysis. **(L)**RAC performed on murine wild-type and APP/PS1 whole brain lysate. **(M)**Cell fractionation of wild-type and APP/PS1 mice brains with immunoprecipitation of mTOR and immunoblot for iNOS, nNOS, eNOS, PIKfyve, VPS34, and mTOR, with loading controls actin and LAMP1. **(N)**Lysosomal NOS and **(O)**proteolysis activity of lysosomes isolated from murine wild-type and APP/PS1 brains (p<0.02). **(P)**RAC performed on HEK293 cells subjected to EBSS-starvation and supplemented with either L-arginine or L-NAME, and **(Q)**its respective densitometry analysis. **(R)** NOS activity of rat brain-derived lysosomes pre-treated with L-arginine, L-NAME or Torin-1, and **(S)** lysosomes derived from ATG5-knockout MEFs. Sample size numbers are expressed in parenthesis, and pooled data is expressed as mean (SEM). *p<0.05;**p<0.01;***p<0.001;Scale bar:10µm. **Supplementary data: Figure 3-figure supplement 1**

### Cys423Ala-mTOR reverses AD pathology

To identify the site of S-nitrosylation on mTOR, we performed a mutant screen based on *in-silico* prediction(*19*) of cysteine residues that were most susceptible to S-nitrosylation, by performing site-directed mutagenesis (Cys>Ala) of 5 cysteine residues. Cys^423^>Ala mTOR mutant(Cys^423A^-mTOR) was the only mutant that reversed NOC18-mediated autophagic inhibition(**Figure 4A**), lysosomal accumulation(**Figure 4B**), and impaired lysosomal proteolysis(**Figure 4C**) without significant change in p70S6K phosphorylation, suggesting mTORC1-independent effects(**Figure 4-figure supplement 1 A-D**). Furthermore, *in silico* modelling suggests no significant protein structure changes with Cys423Ala mutation(**Figure 4-figure supplement 1 E,F**) that could affect its kinase activity. Correspondingly, we observed decreased S-nitrosylation levels with Cys^423A^-mTOR(**Figure 4D**). We next observed that mTOR S-nitrosylation levels were elevated in 3 age-matched pairs of AD patient-derived fibroblasts:AG06869, AG07377, and AG08243, compared with healthy controls ND34791, AG04148, and GM01681 respectively(**Figure 4E,F**), and hypothesized that L-NAME or Cys^423A^-mTOR overexpression could be novel therapeutic strategies in AD. We found that both L-NAME and Cys^423A^-mTOR overexpression reversed lysosomal accumulation(**Figure 4G,H; Figure 4-figure supplement 1 G,H)**, impaired lysosomal proteolysis(**Figure 4I,J; Figure 4-figure supplement 1 I,J**) and autophagy(**Figure 4K,L; Figure 4-figure supplement 1 K,L**) in AD fibroblasts, whereas no significant change in lysosomal proteolysis or autophagic activity was observed in the paired healthy controls. As a proof-of-concept, we next treated APP/PS1 mouse-derived hippocampi with L-NAME, and observed sustained late-LTP, typically absent in untreated APP/PS1 hippocampi(**Figure 4M,N**). Taken together, our data suggests that NOS-mediated mTOR S-nitrosylation is implicated in AD and could be a potential therapeutic target in reversing neurodegenerative processes stemming from autophagic and lysosomal failure.

**Figure 4:**
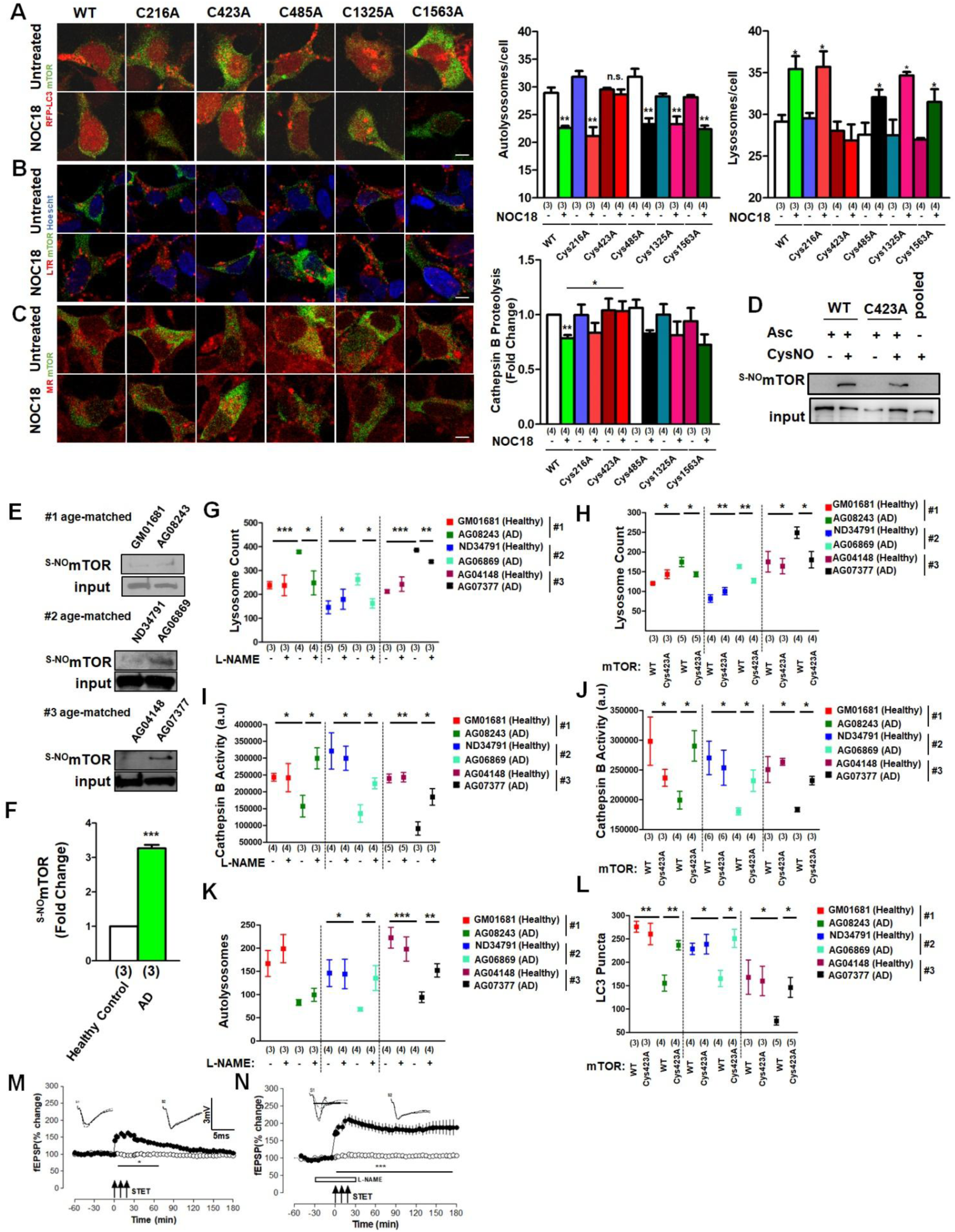
Cys423 S-nitrosylation of mTOR by lysosomal NOS is a therapeutic target in AD. **(A)**Overexpression of LC3, and **(B)**Lysotracker Red and **(C)**Magic Red staining of HEK293 cells transfected with mutant mTOR, and their corresponding quantification graphs. **(D)**RAC performed on wild-type mTOR and Cys^423A^-mTOR, and **(E)**3 pairs of age-matched healthy controls(GM01681, ND34791, AG04148) and AD patient-derived fibroblasts(AG08243, AG06869, AG07377 respectively), with **(F)**its respective densitometry. **(G)**Lysosomal counts of healthy control and AD patient-derived fibroblasts subjected to L-NAME treatment and **(H)**Cys^423A^-mTOR overexpression. (Lines are delineated for easy interpretation of a healthy control with the paired AD patient) **(I)**Lysosomal proteolysis activity of healthy control and AD patient-derived fibroblasts subjected to L-NAME treatment and **(J)**Cys^423A^-mTOR overexpression. **(K)**Autolysosome counts of healthy control and AD patient-derived fibroblasts subjected to L-NAME treatment and (**L)**Cys^423A^-mTOR overexpression. **(M)**L-LTP induced by STET in untreated APP/PS1 mice and **(N)**L-NAME pre-treatment. Filled circles represent S1 recording, while open circles represent stable control input S2. Analog traces represent fEPSP traces recorded from synaptic inputs S1 and S2 30-minutes before (dotted), 30-minutes (broken) and 3-hours (continuous) after tetanisation. Sample size numbers are expressed in parenthesis, and pooled data is expressed as mean (SEM). *p<0.05;**p<0.01;***p<0.001;Scale bar:10µm. **Supplementary data: Figure 4-figure supplement 1**

## Discussion

mTOR has been classically defined as a kinase involved in regulation of the autophagosome-lysosomal pathway downstream of AMPK and PI3K-Akt signalling(*20*), whereby suppression of mTORC1 results in lysosomal and autophagic activation(*21*). Here our study re-defines our understanding of the mTOR pathway with a kinase-independent, non-canonical mechanism of mTOR regulation via regulation of phosphoinositides. Previous studies reported that activation of lysosomal iNOS disrupts lysosomal function(*14*), while lysosomal arginine, a substrate of NOS, plays a critical role in lysosomal and autophagic regulation via mTORC1(*5*). Our study reconciles these observations by proposing that lysosomal arginine serves as a marker of cellular nutrient status via mTOR nitrosylation. Finally, S-nitrosylation of mTOR is elevated in both Alzheimer’s disease mice and human models, while antagonism of NOS activity or overexpression of Cys423Ala-mTOR mutant reversed lysosomal and autophagic deficits in Alzheimer’s disease patient-derived fibroblasts, and LTP deficits in APP/PS1 hippocampi. In summary, our findings are in keeping with a phosphorylation-dominant mTOR signalling pathway in autophagic regulation which is sensitive to mTORC1 inhibitors, and provide an additional layer of complexity of mTOR regulation of autophagy and lysosomes (**Figure 5**). We propose that S-nitrosylation of mTOR is a potential therapeutic target for autophagic and lysosomal activation in aging and neurodegenerative diseases that minimizes systemic side effects of mTORC1 kinase inhibition precluding clinical utility.

**Figure 5:**
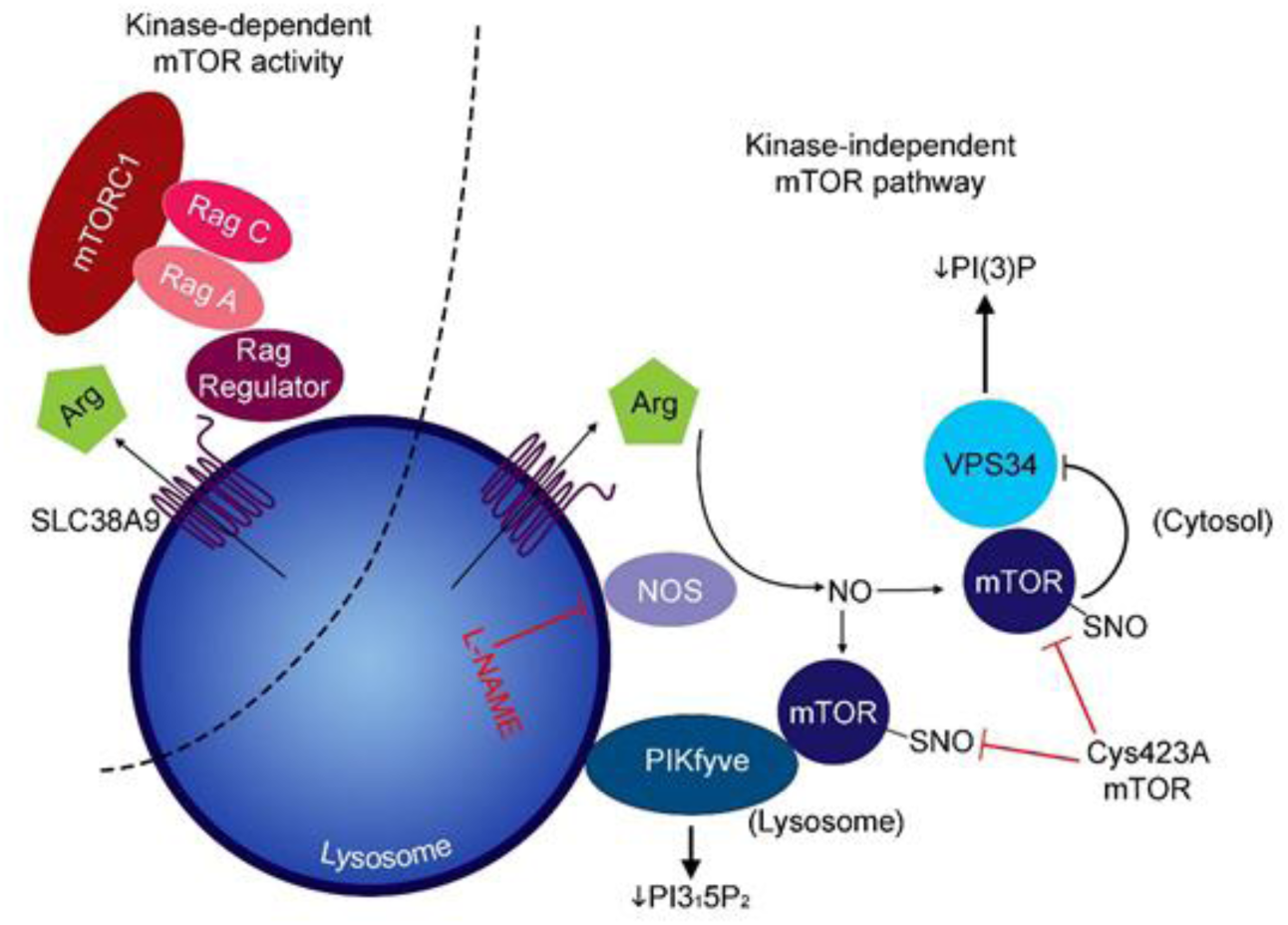
Kinase-independent effects of mTOR S-nitrosylation. Kinase-dependent mTORC1 activity is associated with arginine-efflux lysosomal transporter SLC38A9 described in previous studies. Observations in this study suggest lysosomal arginine, under nutrient-replete conditions, is transported out of the lysosome and converted to NO by lysosomal nitric oxide synthases and consequently nitrosylating mTOR. S-nitrosylated mTOR subsequently interacts with lysosomal PIKfyve and cytosolic VPS34 to decrease PI(3,5)P_2_ and PI(3)P respectively, thus downregulating lysosomal function and autophagy.

## Materials and Methods

### Cell culture

Human embryonic kidney-293 (HEK293) cells were grown in DMEM containing 10% FBS and 5% penicillin/streptomycin in 5% CO_2_. To monitor the effect of NO on autophagy, cells were treated with NO donor NOC-18 (4µM for 24 hr, Santa Cruz), rapamycin (200nM, Sigma), Torin-1 (500nM or 1µM, Cayman Chemicals), wortmannin (200nM, Cayman Chemicals), L-NAME (50µM, Sigma), L-arginine (0.4mM, Sigma), YM201636 (1µM, Cayman Chemicals), PI(3)P-diC16 (1µM, Echelon), PI(4)P-diC16 (1µM, Echelon), PI(5)P-diC16 (1µM, Echelon), and PI(3,5)P_2_-diC8 (1µM, Echelon). HEK293 cells were obtained from ATCC, and grown in DMEM containing 10% FBS and 5% penicillin/streptomycin. ATG5 wild-type and knock-out mouse embryonic fibroblasts were kind gifts from Dr Noboru Mizushima from Tokyo Medical and Dental University. Transfection of plasmids or siRNA were performed at 70% and 50% confluency respectively. siRNAs were transfected to achieve 0.5nM final concentration in media. Lipofectamine 2000 was used for transfection of HEK293 cells, and Lipofectamine 3000 for transfection of patient-derived fibroblast cell lines as per manufacturer instructions for siRNAs and plasmids.

### Animal Models

All the experimental procedures were done in accordance with the guidelines and protocols approved by the National University of Singapore (NUS) Institutional Review Board and according to the guidelines of the Institutional Animal Care & Use Committee (IACUC), NUS, Singapore. The AD mice model bears a chimeric mouse/human APP with mutations linked to familial AD and a human PS1 carrying the exon-9–deleted variant associated with familial AD under control of a prion promoter element (APPSwe/PS1dE). 4 to 9 month old wild-type or APP/PS1 mice were used for biochemical experiments, while age-matched male nontransgenic C57BL/6 mice were used as wild-type controls for electrophysiological studies. A total of 10 APP/PS1 mice, and its corresponding wild-type controls were used for the electrophysiological studies, while an additional 5 APP/PS1 mice and 5 wild-type control mice were used for biochemical characterization. Rat whole brains were obtained from 6-week old wild-type rats, while cortical neurons from wild-type rats were isolated in accordance to the protocol below.

### Patient-derived Fibroblasts

Healthy age-matched controls (AG04148, ND034791, GM01681) and idiopathic Alzheimer’s Disease patients (AG07377, AG08243, AG07377) were obtained from Coriell Institute, and overexpressed with GFP-mTOR or the Cys423Ala mTOR mutant with Lipofectamine 3000.

### Rat Primary Neuron Culture

Primary rat cortical neurons were cultured using a modified protocol as described previously(*22*). Briefly, rat embryonic day 16 (E16) to E18 cortical neurons were cultured on glass coverslips coated with poly-D-lysine in Neurobasal medium with B27 supplement (Invitrogen) containing 1% penicillin/streptomycin and 2mM glutamine. 24h after plating, cultures were treated with 1µM cytarabine for 48h to remove contaminating glia. Cells were used at 17 days *in vitro* (DIV 17) for all calcium imaging experiments.

### Cell Fractionation

Fresh rat or murine brain obtained from wild-type or APP-PS1, or SH-SY5Y cells were equilibrated in lysosomal extraction buffer (135mM K, 45mM Cl, 90mM gluconate, 30mM HEPES-NaOH, 30mM sucrose, pH 7.40) and dounced 40 times using a homogenezier or syringed 40 times using a 27G needle respectively on ice. Cell lysates were then subjected to an initial 8000g spin, followed by a 20000g spin. Lysosomal pellet is collected at the end of the 20000g spin, while the cytosolic fraction is obtained from the supernatant.

### Lysosomal nitric oxide synthase activity assay

Lysosomes are incubated with 1mg/ml L-arginine and in lysosomal extraction buffer for 3 hours, and NO metabolites are measured using the diaminonaphthalene (DAN) assay. Briefly supernatant from the lysosome suspension is added to a final concentration of 1mg/ml DAN dissolved in 0.1M HCl, and incubated at 37°C for 15 minutes. The reaction is neutralized using equimolar volumes of 0.2M NaOH, and the reaction buffer is detected at 360 nm excitation, and 415 nm emission.

### Immunoblotting

The primary antibodies and their respective dilutions used in immunoblotting are as followed, listed in alphabetical order: AKT (pan) (C67E7) (Cell Signalling Technology (CST) #4691, 1:1000), ATG14 (CST #5504, 1:1000), β-actin (Abcam ab8226, 1:10,000), β-tubulin (E7) (Developmental Studies Hybridoma Bank, 1:2000), Beclin-1 (CST #3495, 1:1000), GFP (Roche Cat No 11814460001, 1:3000), mTOR (CST #2972, 1:1000), Rictor (CST #2114, 1:500), Raptor (CST #2280, 1:500), p70/S6 kinase (CST #2708, 1:1000), phospho-Akt^T308^ (CST #9275, 1:1000), phospho-Akt^S473^ (CST #4060, 1:1000), phospho-mTOR^S2448^ (CST #2974, 1:1000), phospho-p70/S6 kinase^T389^ (CST #9204, 1:1000), phospho-ULK1^S555^ (D1H4) (CST #5869, 1:1000), phospho-ULK1^S757^ (CST #14202, 1:1000), PIKfyve (Santa Cruz, sc-100408 1:1000), ULK1 (D8H5) (CST #8054, 1:1000), and VPS34 (CST #4263, 1:1000).

### Immunoprecipitation

1ug of mTOR, ATG14, Beclin-1, and VPS34 antibody was added into cell lysates, and incubated overnight with Dynabeads. Immunoprecipitates were washed extensively with cold PBS, and eluted with 6x SDS-loading buffer.

### Resin-Assisted Capture

Detection of S-nitrosylation of mTOR was done with resin-assisted capture method as previously described(*23*).

### Plasmids

GFP-Rab7A^WT^ and Rab7A^Q67L^ mutant were gifts from Dr Gia Voeltz (Addgene plasmid #61803)(*24*), GFP-LAMP1 was a gift from Dr Ron Vale (Addgene plasmid #16290)(*25*), GFP-EEA1 was a gift from Dr Silvia Corvera (Addgene plasmid #42307)(*26*), and RFP-LC3 was a gift from Dr Tamotsu Yoshimori (Addgene plasmid #21075)(*27*). GFP-mTOR cysteine mutants – Cys216Ala, Cys423Ala, Cys485Ala, Cys1325Ala and Cys1563Ala were generated by site-directed mutagenesis.

### Generation of GFP-TFEB SH-SY5Y stable cell line

GFP-TFEB expression construct and blasticidin-resistant vector were purchased from the Addgene plasmid repository. SY5Y cells were co-transfected with 1.0μg of GFP-TFEB construct together with 1.0μg of blasticidin-resistant vector for 72h. Following transfection, positive cells co-expressing GFP-TFEB and blasticidin-resistant vector were selected by killing with 10μg/ml blasticidin (Invitrogen) for 5-7 days. Co-transfected cells that survived blasticidin treatment were typsinized and re-seeded to allow growth of single cell colonies under the influence of 10μg/ml blasticidin. Positive clones were isolated, and allowed to grow and expand in the presence of 2μg/ml blasticidin. The GFP-TFEB stable SY5Y cells were subsequently maintained in serum-supplemented media supplemented with 2μg/ml blasticidin.

### Confocal Microscopy

Cells were stained with Lysotracker Red (1:2000, Thermoscientific) for 1 hr, Cathepsin B substrate Magic Red (1:600, Immunochemistry) for 1 hr before fixing in 2.5% PFA for 20 minutes. For analysis of autophagosomes or autolysosomes, cells were transfected with RFP-LC3 or tandem-fluorescent LC3, and subjected to fixing in 2.5% PFA for 20 minutes after treatment. Cells were extensively washed and stained with Hoescht 34222 (1:500, Immunochemistry). Confocal images were obtained from Zeiss LSM510 confocal microscope. Magic Red intensity was obtained using the AlexaFluor-594 excitation-emission setting, with detection gain set between 750-850, and all images within each biological replicate were taken with the same settings. Lysotracker Confocal images were analysed using a semi-automated puncta counter using ImageJ for lysosomes and LC3 puncta, while densitometry-based analysis was performed for Magic Red fluorescent intensity.

### Lysosomal enzyme activity measurement

Lysosomes extracted as described above were resuspended in 1µM 4-methylumbelliferyl β-D-glucopyranoside (Sigma) for measurement of β-glucosidase activity, or 1µM 4-methylumbelliferyl β-D-galactopyranoside (Sigma) for measurement of β-galactosidase activity. To analyse effects of lysosomal calcium on enzymatic activity, lysosomes were dialyzed with the MES standard buffers as described above, and incubated with 20µM BAPTA-AM. Enzymatic reaction is quenched with 10Mm NaOH, and fluorescence intensities of excitation 360nm and emission of 450nm were recorded.

### Hippocampal Slice Preparation and Electrophysiology

Hippocampal slices were prepared from 4-5 month old APP/PS1 mice. Mice were anaesthetized using CO_2_ and then decapitated immediately, after which the brains were quickly removed and cooled in 4-6 °C artificial cerebrospinal fluid (ACSF). Hippocampi were dissected, and transverse hippocampal slices (400 µm) were sectioned by using a manual tissue chopper. Slices were then immediately incubated in an interface chamber (Scientific System Design) maintained at 32 °C for 2–3 h, continuously perfused with oxygenated ACSF at a flow rate of 0.80 mL/min. The ACSF contained the following (in mM): 124 NaCl, 4.9 KCl, 1.2 KH_2_PO_4_, 2.0 MgSO_4_, 2.0 CaCl_2_, 24.6 NaHCO_3_, and 10 D-glucose, equilibrated with 95% O_2_/5% CO_2_ (32 L/h). In all electrophysiological recordings, two-pathway experiments were performed, i.e., two monopolar lacquer-coated, stainless-steel electrodes (5 MΩ; AM Systems) were positioned at an adequate distance within the stratum radiatum of the CA1 region for stimulating two independent synaptic inputs S1 and S2 of a neuronal population, thus evoking field EPSPs (fEPSP) from Schaffer collateral/commissural-CA1 synapses. For recording the fEPSP, (measured as its initial slope function), one electrode (5 MΩ; AM Systems) was placed in the CA1 apical dendritic layer, and signals were amplified by a differential amplifier (model 1700; AM Systems). The signals were digitized by using a CED 1401 analog-to-digital converter (Cambridge Electronic Design).

After the preincubation period, a synaptic input-output curve (afferent stimulation vs. fEPSP slope) was generated. Test stimulation intensity was adjusted to elicit fEPSP slope of 40% of the maximal EPSP response for synaptic inputs S1 and S2. For L-LTP induction, (a strong tetanisation) STET protocol involving repeated high-frequency stimulation (three bursts of 100 pulses for 1 s (100 Hz) every 10 min) was performed. The pulse width was 0.2 ms per phase and had double length in comparison with test pulse width. The slopes of the fEPSPs were monitored online. Four 0.2-Hz biphasic constant-current pulses (0.1 ms per polarity) were used for baseline recording at each time point.

The average values of the slope function of the field excitatory postsynaptic potential (fEPSP) per time point were analyzed using Wilcoxon signed-rank test when compared within the group or the Mann–Whitney *U* test when compared between groups.

### Phosphoinositide profiling

Phosphoinositide extraction was adapted from Kiefer *et al* (*12*) and Nasuhoglu *et al (28)* as previously described. Chloroform (4 mL) was added to the cell pellet and agitated for 10mins followed by addition of 2M HCl (2 mL), 2xPhosSTOP (2 mL), 2M NaCl (0.3 mL) and Methanol (4 mL) which contain 3mM BHT, 1mM NaF and 1xPhosSTOP. The mixture was vigorously vortexed and centrifuged. Lower organic phase was separated and a second extraction with chloroform (4 mL) was conducted. The combined organic phase was dried under a stream of nitrogen. Deacylation was then carried out by addition of 4ml of MeNH_2_ with the following composition water/methanol/n-butanol (43:46:11) at 55°C for 1 hr. The solvent was evaporated under reduced pressure and extracted with water and an organic mixture of ethyl formate/n-butanol/petroleum ether (1/20/4). The aqueous phase was concentrated and dry under vacuum. Freeze-dried samples were added with 45 μL of water and 5 μL of 2-(4-hydroxyphenylazo)benzoic acid at 1000ppm as internal standard. Samples were vortexed vigorously and centrifuged at 6000 rpm for 10mins, before sonication was performed for 15mins. This was followed by vigorous vortexing and centrifugation before injecting into the high-performance liquid chromatography (HPLC) system. HPLC separation was performed on an Agilent 1100 coupled with LCQ Fleet Ion trap Mass spectrometer. An Agilent Zorbax Eclipse Plus C18 (2.1×150mm) column was used for the separation.

HPLC mobile phase A: water containing 5mM N,N-dimethylhexylamine and 4mM glacial acetic acid. Mobile phase B: methanol containing 5mM N,N-dimethylhexylamine and 4mM glacial acetic acid. Solvent gradient, 0 min 15% B, 60 min 50% B, 65 min 100% B and remain 100% B for 15 min. The HPLC condition is modified from the report of Peter Küenzi et al(*29*). Spectra was recorded from m/z of 200 to 600 in the negative mode. The chromatogram obtained were smoothed and area under the respective phosphoinositide peaks were integrated and normalized with PI.

### *In silico* Protein Modelling

*In silico* modelling of the protein structures of wild-type mTOR and Cys423Ala-mTOR mutant is done using the SWISS-MODEL open-access software as previously described(*30*).

## Statistical Analysis

2-way Student’s t-test was used for analysis of difference of means for lysosomal count, autolysosomal and autophagosome counts, lysosomal proteolysis indices, lysosomal perinuclear distribution, and Western blot densitometry. Paired Student’s t-test was used for analysis of lysosomal proteolytic activity in APP-PS1 and wild-type mice brains. All graphs are represented as mean ± SEM, and the number of independent biological replicates or sample size are indicated in parenthesis. A p-value of less than 0.05 is considered statistically significant.

## Acknowledgements

We would like to thank Dr. Noboru Mizushima for the ATG5-knockout MEF cell line. We would like to thank the patients who have contributed their samples for this research study.

## References

1. A. H. Tsang, Y. I. Lee, H. S. Ko, J. M. Savitt, O. Pletnikova, J. C. Troncoso, V. L. Dawson, T. M. Dawson, K. K. Chung, S-nitrosylation of XIAP compromises neuronal survival in Parkinson’s disease. Proc Natl Acad Sci U S A 106, 4900–4905 (2009).

2. M. E. Conway, M. Harris, S-nitrosylation of the thioredoxin-like domains of protein disulfide isomerase and its role in neurodegenerative conditions. Front Chem 3, 27 (2015).

3. A. Martinez-Ruiz, S. Lamas, S-nitrosylation: a potential new paradigm in signal transduction. Cardiovasc Res 62, 43–52 (2004).

4. K. K. Chung, B. Thomas, X. Li, O. Pletnikova, J. C. Troncoso, L. Marsh, V. L. Dawson, T. M. Dawson, S-nitrosylation of parkin regulates ubiquitination and compromises parkin’s protective function. Science 304, 1328–1331 (2004).

5. G. A. Wyant, M. Abu-Remaileh, R. L. Wolfson, W. W. Chen, E. Freinkman, L. V. Danai, M. G. Vander Heiden, D. M. Sabatini, mTORC1 Activator SLC38A9 Is Required to Efflux Essential Amino Acids from Lysosomes and Use Protein as a Nutrient. Cell 171, 642–654 e612 (2017).

6. S. Wang, Z. Y. Tsun, R. L. Wolfson, K. Shen, G. A. Wyant, M. E. Plovanich, E. D. Yuan, T. D. Jones, L. Chantranupong, W. Comb, T. Wang, L. Bar-Peled, R. Zoncu, C. Straub, C. Kim, J. Park, B. L. Sabatini, D. M. Sabatini, Metabolism. Lysosomal amino acid transporter SLC38A9 signals arginine sufficiency to mTORC1. Science 347, 188–194 (2015).

7. S. Zahid, R. Khan, M. Oellerich, N. Ahmed, A. R. Asif, Differential S-nitrosylation of proteins in Alzheimer’s disease. Neuroscience 256, 126–136 (2014).

8. T. S. Wijasa, M. Sylvester, N. Brocke-Ahmadinejad, S. Schwartz, F. Santarelli, V. Gieselmann, T. Klockgether, F. Brosseron, M. T. Heneka, Quantitative proteomics of synaptosome S-nitrosylation in Alzheimer’s disease. J Neurochem 152, 710–726 (2020).

9. P. Liu, M. S. Fleete, Y. Jing, N. D. Collie, M. A. Curtis, H. J. Waldvogel, R. L. Faull, W. C. Abraham, H. Zhang, Altered arginine metabolism in Alzheimer’s disease brains. Neurobiol Aging 35, 1992–2003 (2014).

10. D. H. Bergin, Y. Jing, B. G. Mockett, H. Zhang, W. C. Abraham, P. Liu, Altered plasma arginine metabolome precedes behavioural and brain arginine metabolomic profile changes in the APPswe/PS1DeltaE9 mouse model of Alzheimer’s disease. Transl Psychiatry 8, 108 (2018).

11. E. Barbero-Camps, V. Roca-Agujetas, I. Bartolessis, C. de Dios, J. C. Fernandez-Checa, M. Mari, A. Morales, T. Hartmann, A. Colell, Cholesterol impairs autophagy-mediated clearance of amyloid beta while promoting its secretion. Autophagy 14, 1129–1154 (2018).

12. Y. Wang, E. Mandelkow, Degradation of tau protein by autophagy and proteasomal pathways. Biochem Soc Trans 40, 644–652 (2012).

13. D. S. Yang, P. Stavrides, P. S. Mohan, S. Kaushik, A. Kumar, M. Ohno, S. D. Schmidt, D. Wesson, U. Bandyopadhyay, Y. Jiang, M. Pawlik, C. M. Peterhoff, A. J. Yang, D. A. Wilson, P. St George-Hyslop, D. Westaway, P. M. Mathews, E. Levy, A. M. Cuervo, R. A. Nixon, Reversal of autophagy dysfunction in the TgCRND8 mouse model of Alzheimer’s disease ameliorates amyloid pathologies and memory deficits. Brain 134, 258–277 (2011).

14. Q. Qian, Z. Zhang, M. Li, K. Savage, D. Cheng, A. J. Rauckhorst, J. A. Ankrum, E. B. Taylor, W. X. Ding, Y. Xiao, H. J. Cao, L. Yang, Hepatic Lysosomal iNOS Activity Impairs Autophagy in Obesity. Cell Mol Gastroenterol Hepatol 8, 95–110 (2019).

15. S. Sarkar, V. I. Korolchuk, M. Renna, S. Imarisio, A. Fleming, A. Williams, M. Garcia-Arencibia, C. Rose, S. Luo, B. R. Underwood, G. Kroemer, C. J. O’Kane, D. C. Rubinsztein, Complex inhibitory effects of nitric oxide on autophagy. Mol Cell 43, 19–32 (2011).

16. H. X. Yuan, R. C. Russell, K. L. Guan, Regulation of PIK3C3/VPS34 complexes by MTOR in nutrient stress-induced autophagy. Autophagy 9, 1983–1995 (2013).

17. C. H. Choy, G. Saffi, M. A. Gray, C. Wallace, R. M. Dayam, Z. A. Ou, G. Lenk, R. Puertollano, S. C. Watkins, R. J. Botelho, Lysosome enlargement during inhibition of the lipid kinase PIKfyve proceeds through lysosome coalescence. J Cell Sci 131, (2018).

18. M. Vicinanza, V. I. Korolchuk, A. Ashkenazi, C. Puri, F. M. Menzies, J. H. Clarke, D. C. Rubinsztein, PI(5)P regulates autophagosome biogenesis. Mol Cell 57, 219–234 (2015).

19. Y. Xue, Z. Liu, X. Gao, C. Jin, L. Wen, X. Yao, J. Ren, GPS-SNO: computational prediction of protein S-nitrosylation sites with a modified GPS algorithm. PLoS One 5, e11290 (2010).

20. K. Kawauchi, T. Ogasawara, M. Yasuyama, K. Otsuka, O. Yamada, Regulation and importance of the PI3K/Akt/mTOR signaling pathway in hematologic malignancies. Anticancer Agents Med Chem 9, 1024–1038 (2009).

21. R. Puertollano, mTOR and lysosome regulation. F1000Prime Rep 6, 52 (2014).

22. C. P. Poore, J. R. Sundaram, T. K. Pareek, A. Fu, N. Amin, N. E. Mohamed, Y. L. Zheng, A. X. Goh, M. K. Lai, N. Y. Ip, H. C. Pant, S. Kesavapany, Cdk5-mediated phosphorylation of delta-catenin regulates its localization and GluR2-mediated synaptic activity. J Neurosci 30, 8457–8467 (2010).

23. M. T. Forrester, J. W. Thompson, M. W. Foster, L. Nogueira, M. A. Moseley, J. S. Stamler, Proteomic analysis of S-nitrosylation and denitrosylation by resin-assisted capture. Nat Biotechnol 27, 557–559 (2009).

24. A. A. Rowland, P. J. Chitwood, M. J. Phillips, G. K. Voeltz, ER contact sites define the position and timing of endosome fission. Cell 159, 1027–1041 (2014).

25. A. A. Minin, A. V. Kulik, F. K. Gyoeva, Y. Li, G. Goshima, V. I. Gelfand, Regulation of mitochondria distribution by RhoA and formins. J Cell Sci 119, 659–670 (2006).

26. D. C. Lawe, V. Patki, R. Heller-Harrison, D. Lambright, S. Corvera, The FYVE domain of early endosome antigen 1 is required for both phosphatidylinositol 3-phosphate and Rab5 binding. Critical role of this dual interaction for endosomal localization. J Biol Chem 275, 3699–3705 (2000).

27. S. Kimura, T. Noda, T. Yoshimori, Dissection of the autophagosome maturation process by a novel reporter protein, tandem fluorescent-tagged LC3. Autophagy 3, 452–460 (2007).

28. S. Kiefer, J. Rogger, A. Melone, A. C. Mertz, A. Koryakina, M. Hamburger, P. Kuenzi, Separation and detection of all phosphoinositide isomers by ESI-MS. J Pharm Biomed Anal 53, 552–558 (2010).

29. C. Nasuhoglu, S. Feng, J. Mao, M. Yamamoto, H. L. Yin, S. Earnest, B. Barylko, J. P. Albanesi, D. W. Hilgemann, Nonradioactive analysis of phosphatidylinositides and other anionic phospholipids by anion-exchange high-performance liquid chromatography with suppressed conductivity detection. Anal Biochem 301, 243–254 (2002).

30. A. Waterhouse, M. Bertoni, S. Bienert, G. Studer, G. Tauriello, R. Gumienny, F. T. Heer, T. A. P. de Beer, C. Rempfer, L. Bordoli, R. Lepore, T. Schwede, SWISS-MODEL: homology modelling of protein structures and complexes. Nucleic Acids Res 46, W296–W303 (2018).

