## Supplementary Information for "mTOR S-nitrosylation inhibits autophagy and lysosomal proteolysis"

**Page 2-3: Figure 1-figure supplement 1: NO regulates autophagy and lysosomal proteolysis in an mTORC1-independent mechanism.**

**Page 4-5: Figure 2-figure supplement 1: NO-mediated phosphoinositide depletion mediates lysosomal and autophagic impairment.**

**Page 6: Figure 3-figure supplement 1: L-Arginine regulates lysosomal and autophagosomal biogenesis, but not proteolysis under physiological conditions.**

**Page 7-8: Figure 4-figure supplement 1: Cys423Ala (C423A)-mTOR mutant reverses lysosomal and autophagic deficits in idiopathic AD patient-derived fibroblasts.**

### Figure 1-figure supplement 1

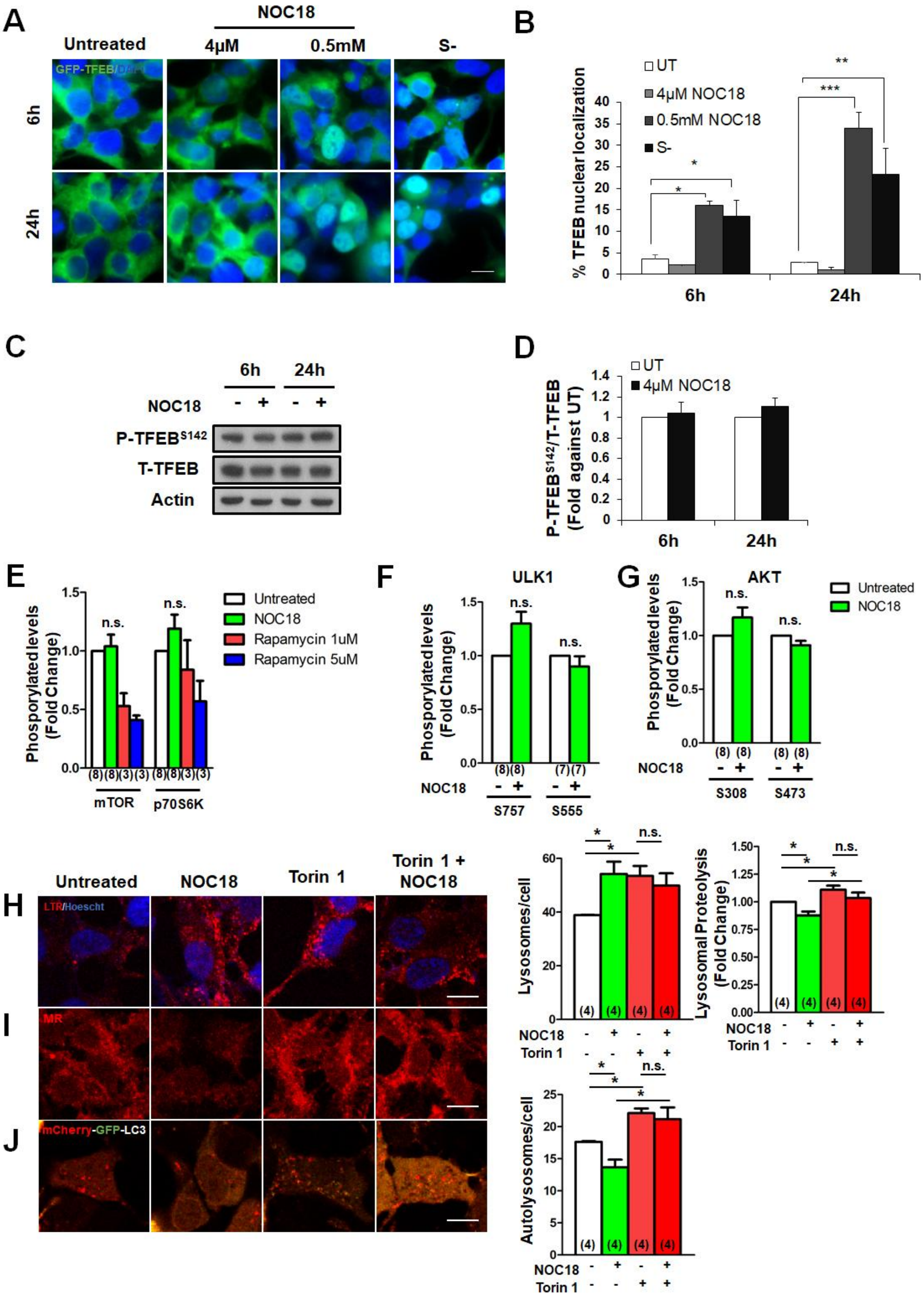

**Figure 1-figure supplement 1**: NO regulates autophagy and lysosomal proteolysis in an mTORC1-independent mechanism. **(A)** GFP-TFEB localization in GFP-TFEB stably-expressing SH-SY5Y cells, with serum starvation (S-), and low- and high-dose NOC18 treatment. **(B)** Quantified TFEB nuclear localization (*$p<0.05$; **$p<0.01$; ***$p<0.001$). **(C)** Phosphorylation levels of TFEB at serine 142 with NOC18 treatment, and **(D)** its corresponding densitometry analysis. Densitometry analysis of **(E)** phosphorylated-mTOR, p70S6K, **(F)** ULK1 and **(G)** AKT. **(H)** Lysotracker Red, **(I)** Magic Red, and **(J)** mCherry-GFP-LC3 overexpression of SH-SY5Y cells treated with Torin-1 and NOC18, with its respective quantification. Numbers in parenthesis are indicative of sample size number. All graphs are expressed as mean ± SEM. Scale bar: 5µm

### Figure 2-figure supplement 1

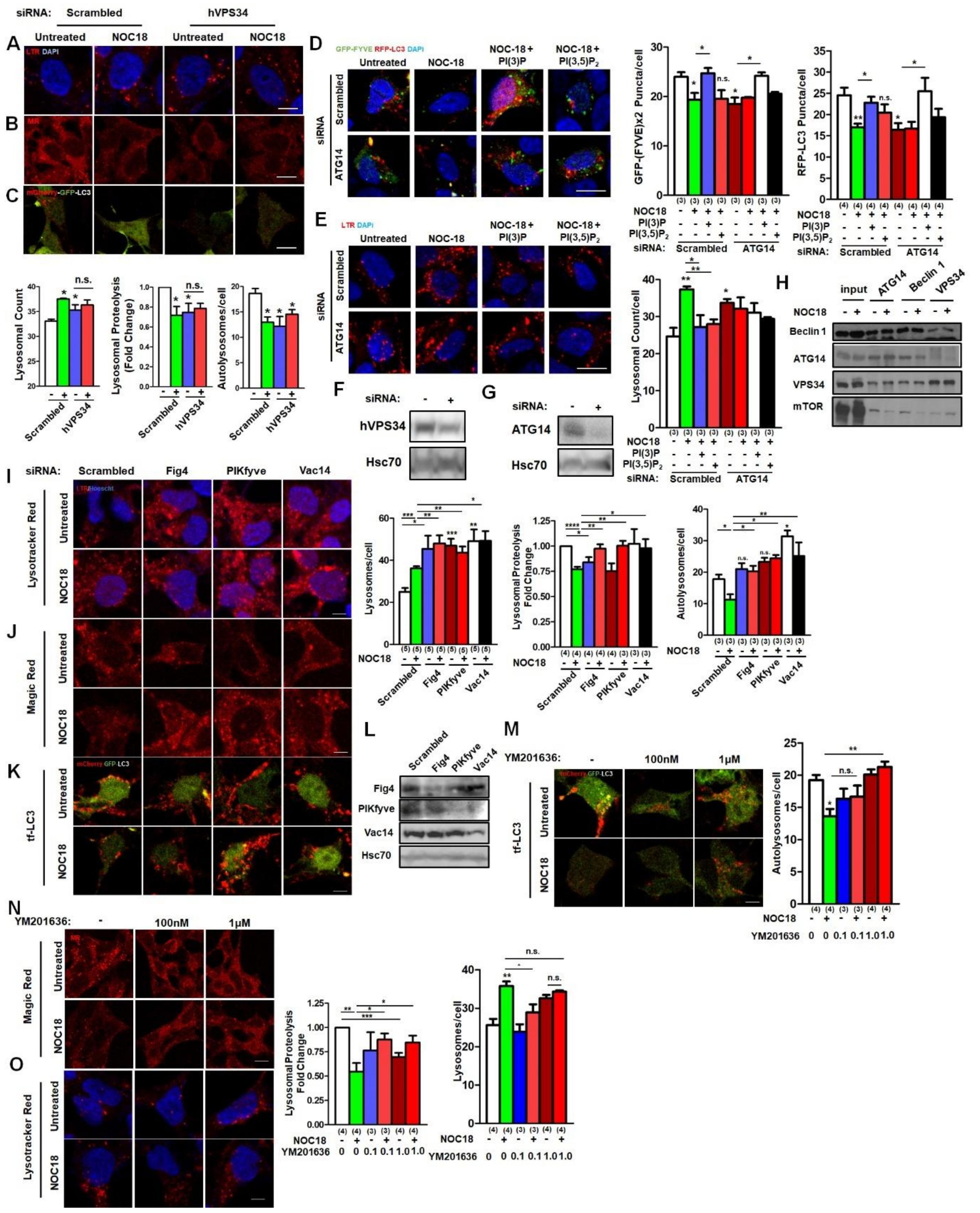


**Figure 2-figure supplement 1:** NO-mediated phosphoinositide depletion mediates lysosomal and autophagic impairment. **(A)** Lysotracker Red, **(B)** Magic Red Cathepsin B staining, and **(C)** mCherry-GFP-LC3 transfection of SH-SY5Y cells with co-transfection with Scrambled or hVPS34 siRNA with their respective quantification of confocal imaging. **(D)** RFP-LC3 and GFP-(FYVE)x2 co-expression and **(E)** Lysotracker Red staining in scrambled or ATG14 siRNA-knockdown SH-SY5Y cells treated with NOC-18, PI(3)P or PI(3,5)$P_{2.}$ **(F)** Immunoblot of hVPS34 and **(G)** ATG14 siRNA knockdown efficiencies. **(H)** Immunoprecipitation of ATG14, Beclin-1 and VPS34. **(I)** Lysotracker Red and **(J)** Magic Red Cathepsin B staining, and **(K)** co-transfection of mCherry-GFP-LC3 (tf-LC3) in SH-SY5Y cells transfected with scrambled, Fig4-, PIKfyve- or Vac14-siRNA, and their respective quantification. **(L)** Knockdown efficacy of Fig4, PIKfyve and Vac14 siRNA transfection. **(M)** Treatment of YM201636 PIKfyve inhibitor to SH-SY5Y overexpressing tf-LC3 and its respective autolysosome quantification. **(N)** Magic Red staining of SH-SY5Y cells treated with YM201636, and its respective intensity quantification. **(O)** Lysotracker Red staining of SH-SY5Y cells treated with YM201636, and its respective punctum quantification. Numbers in parenthesis are indicative of sample size number; *$p<0.05$;**$p<0.01$;Scale bar: 10µm

**Figure 3-figure supplement 1**

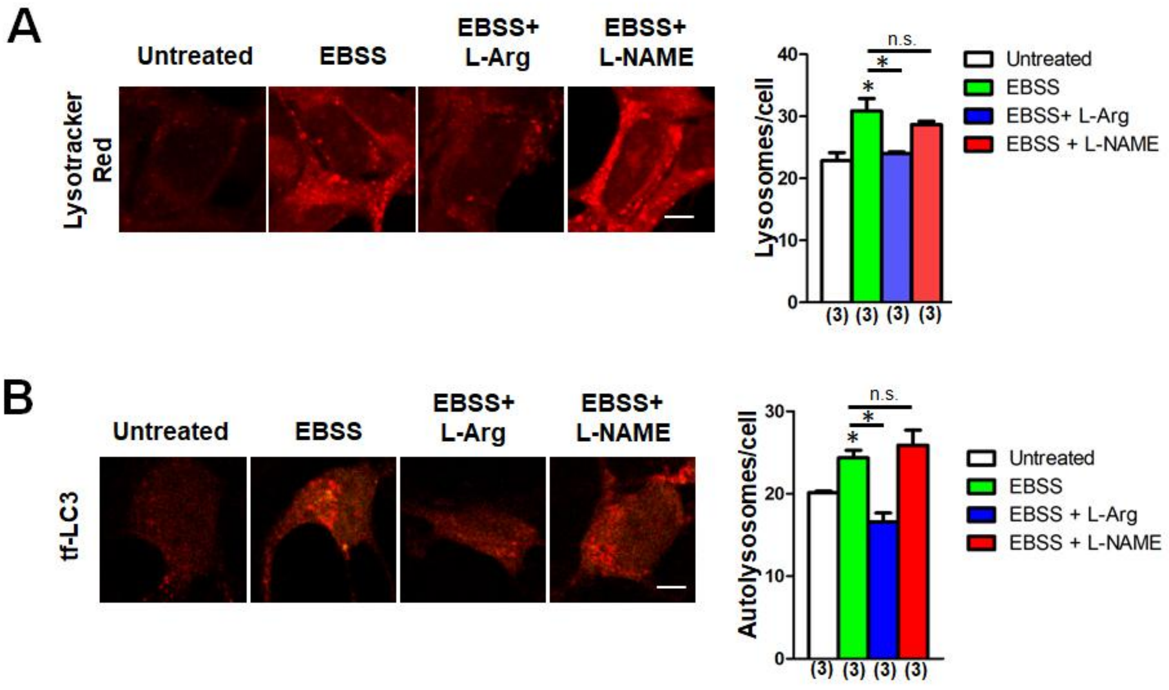


**Figure 3-figure supplement 1:** L-Arginine regulates lysosomal and autophagosomal biogenesis, but not proteolysis under physiological conditions. **(A)** Lysotracker Red and **(B)** tf-LC3 overexpression, and their respective lysosome and autolysosome quantification of SH-SY5Y cells subjected to EBSS starvation with L-Arginine or L-NAME co-treatment. Numbers in parenthesis are indicative of sample size number; *$p<0.05$;Scale bar: 10µm

### Figure 4-figure supplement 1

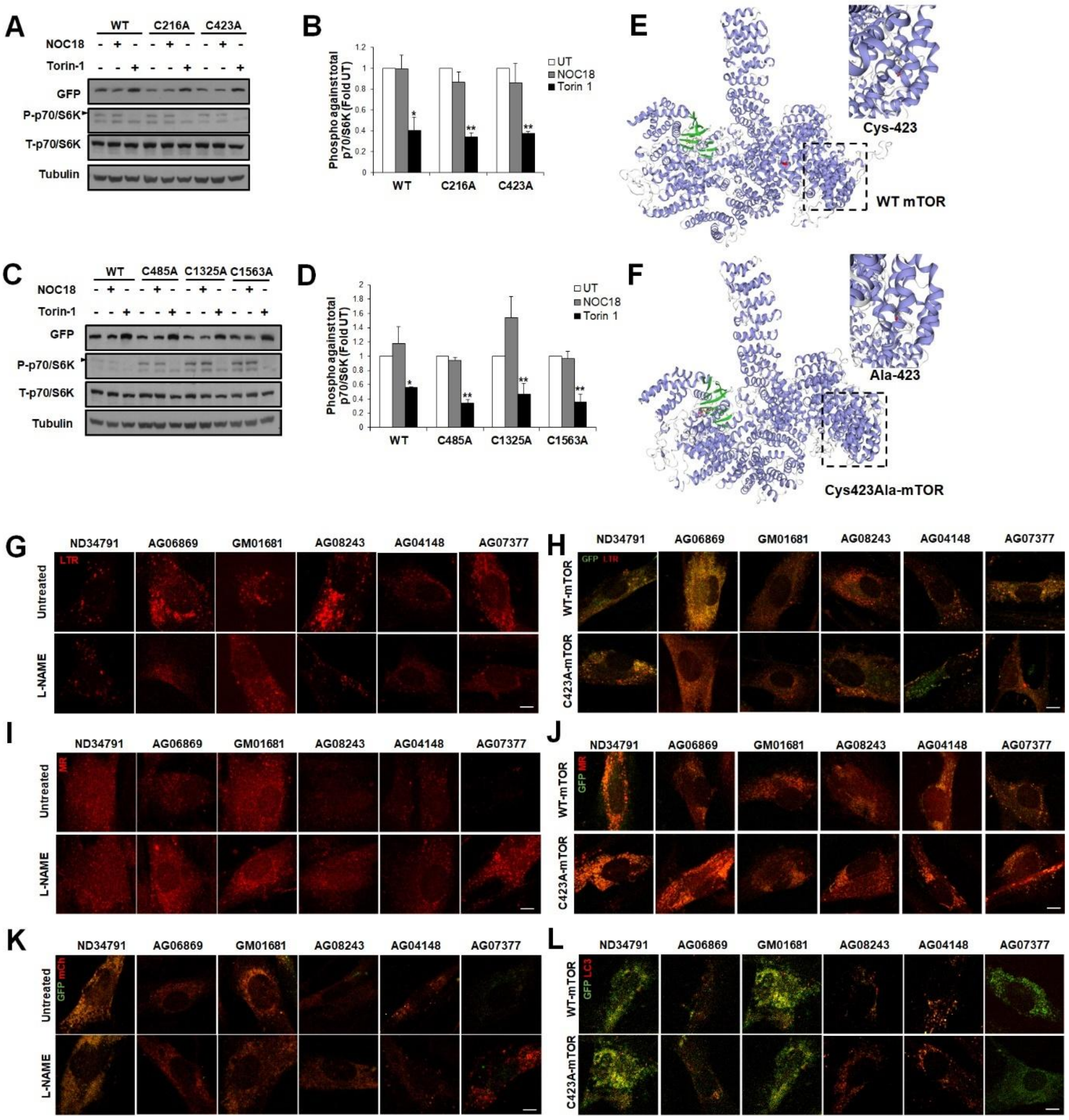


**Figure 4-figure supplement 1:** Cys423Ala (C423A)-mTOR mutant reverses lysosomal and autophagic deficits in idiopathic AD patient-derived fibroblasts. **(A)** p70S6K immunoblot of wild-type, Cys216Ala, and Cys423Ala mTOR mutants and **(B)** its corresponding densitometry analysis. **(C)** p70S6K immunoblot of Cys485Ala, Cys1325Ala, and Cys1563Ala mTOR mutants and **(D)** its corresponding densitometry analysis. In silico modelling of **(E)** wild-type mTOR, and **(F)** mutant Cys423Ala mTOR mutant. Representative pictures of **(G)** Lysotracker-stained patient-derived fibroblast treated with L-NAME or **(H)** transfected with C423A-mTOR**; (I)** Magic Red staining of patient-derived fibroblast treated with L-NAME or **(J)** transfected with C423A-mTOR. **(K)** Overexpression of tf-LC3 in patient-derived fibroblast treated with L-NAME, and **(L)** co-expression of wild type or C423A mTOR with RFP-LC3. *$p<0.05$, **$p<0.01$; Scale-bar: 10μm
